# Uncertainty-aware synthetic lethality prediction with pretrained foundation models

**DOI:** 10.64898/2026.02.25.708096

**Authors:** Kailey Hua, Ellie Haber, Jian Ma

## Abstract

Synthetic lethality (SL) offers a promising paradigm for targeted cancer therapy, yet experimental identification of SL gene pairs remains costly, context-dependent, and biased toward well-studied genes. Existing computational approaches often rely on curated protein-protein interaction (PPI) networks and Gene Ontology (GO) annotations, which limit their ability to generalize to novel genes. Here we introduce Cilantro-sl, a two-stage, graph-free framework that leverages pretrained biological foundation models to predict SL pairs with calibrated uncertainty. In Stage 1, we apply a pretrained single-cell foundation model to bulk RNA-seq profiles of cancer cell lines to obtain context-aware embeddings and perform in silico gene knockouts to generate delta embeddings. These perturbation signals are further conditioned on a data-driven gene prior and supervised with CRISPR viability readouts to learn knockout-aware viability embeddings. In Stage 2, we derive pairwise features from these embeddings and train a lightweight classifier to distinguish SL from non-SL pairs. To enable reliable experimental prioritization, Cilantro-sl incorporates conformal prediction, producing calibrated and interpretable prediction sets that highlight high-confidence SL candidates. Across two evaluation settings, including zero-shot generalization to unseen gene pairs and to unseen genes, ablation analyses show that viability pretraining and the gene prior substantially improve performance while avoiding reliance on PPI and GO features. Cilantro-sl therefore transforms pretrained biological representations into practical, uncertainty-aware hypotheses that support robust and scalable discovery of therapeutic targets.

## 1 Introduction

Synthetic lethal (SL) gene pairs represent a unique genetic relationship in which the simultaneous loss-of-function of two genes leads to cell death, while the loss of either gene alone does not compromise cell survival [1]. SL interactions offer a powerful paradigm for precision oncology: if a tumor harbors a loss-of-function in one gene, therapeutically inhibiting its SL partner can selectively kill cancer cells while sparing normal cells [2]. However, the combinatorial search space is enormous – on the order of 200 million human gene pairs – and true SL interactions are both rare and context dependent, making exhaustive experimental screening infeasible and expensive [3].

Computational approaches have consequently emerged to prioritize SL candidates, including statistical methods (e.g., DAISY [4]), graph-based models (DDGCN [5], KG4SL [6], MVGCN-iSL [7], SLM-GAE [8]), matrix factorization approaches (SL^2^MF [9], GRSMF [10]), and pretrained embedding–based models such as ESM4SL [11]. Despite architectural differences, many rely on curated biological re-sources – including protein-protein interaction (PPI) networks, Gene Ontology (GO) similarity, and SL knowledge graphs – as primary inductive biases. While such priors encode valuable biological structure, they are incomplete and unevenly annotated, potentially limiting generalization to sparsely characterized genes and novel cellular contexts. These limitations motivate complementary approaches that learn gene dependencies directly from perturbation and expression data, rather than predefined interaction topology.

Recent single-cell foundation models (scFMs), such as Geneformer [12] and scGPT [13], learn context-aware cell and gene representations directly from large-scale single-cell RNA sequencing (scRNA-seq) data, capturing functional relationships without explicit PPI or GO supervision. In parallel, pre-trained gene-level embeddings – including protein language models (ESM2 [14]), text-derived gene embeddings (GenePT [15]), and co-expression embeddings (Gene2vec [16]) – encode additional functional signals and expand coverage to sparsely annotated genes, facilitating zero-shot generalization. Together, these pretrained representations enable SL prediction to be formulated as a perturbation-aware representation learning problem, reducing dependence on predefined interaction topology and dense curated annotations.

Another critical barrier is actionability. SL labels are sparse and noisy, and the number of untested gene pairs vastly exceeds labeled examples. In practice, SL models are best suited as *screening tools* to prioritize candidates for costly experiments. Effective prioritization therefore necessitates uncertainty-aware predictions that can rank candidates by confidence and identify interactions with a high likelihood of synthetic lethality. Moreover, because SL interactions are strongly context dependent, meaning a gene pair may be lethal in one setting and neutral in another, models that produce global, cell-line agnostic labels risk overgeneralization. Reliable uncertainty estimates are thus essential both for experimental triage and for accelerating biological discovery [17].

Here, we introduce Cilantro-sl, a framework that reduces reliance on curated interaction networks by leveraging pretrained biological representations and couples SL predictions with rigorous uncertainty quantification. Cilantro-sl fuses scFM-derived cell embeddings with gene priors to extend coverage to understudied genes. We pretrain gene-level viability embeddings on large-scale DepMap CRISPR data to ground representations in perturbation effects and model pairwise SL interactions using these perturbation-aware representations, integrating gene and cell context through lightweight FiLM conditioning [18]. Finally, Cilantro-sl applies conformal prediction to produce calibrated, per-pair confidence sets with finite-sample error control, enabling ranked prioritization of high confidence SL candidates suited for experimental follow-up. Together, these components establish a scalable framework grounded in pretrained biological representations that supports uncertainty-aware, zero-shot SL prediction.

## 2 Methods

### 2.1 Overview of C**ilantro-sl**

Cilantro-sl is a two-stage framework for predicting SL gene pairs that decouples viability-aware single-gene KO representation learning from pairwise SL classification with conformal calibration for rigorous uncertainty quantification. This separation allows the model to first learn transferable perturbation-aware representations and then reuse them flexibly for generalization tasks.

In the first stage, for each cell line and query gene, a Geneformer cell-line embedding is conditioned on a gene embedding **e**_*g*_ through a feature-wise linear modulation (FiLM) layer. The encoder is trained on DepMap CRISPR single-KO viability using a regression objective, learning how perturbing a given gene affects viability in a specific molecular background. Crucially, this stage relies only on pretrained biological representations and observed KO viability effects, and produces a *viability embedding* that encodes gene-specific perturbation effects across diverse cancer cell lines.

In the second stage, for any candidate gene pair, Cilantro-sl constructs pairwise features from the corresponding viability embeddings and trains a binary classifier to distinguish SL from non-SL using curated SL labels. Classifier scores are then converted to calibrated outputs via conformal prediction on a held-out calibration set, yielding coverage guarantees for model confidence and enabling prioritized selection of high-confidence SL candidates for experimental testing.

### 2.2 Pretraining for single KO viability embeddings

We utilize a scFM to construct single-KO viability embeddings, providing context-aware representations of perturbation effects. Unlike models that must learn gene and cell representations from scratch, scFMs provide pretrained gene and cell embeddings that can be readily adapted for downstream tasks such as gene KO viability prediction.

Geneformer [12] is a scFM pretrained on 30 million transcriptomes and is capable of deriving gene and cell representations in a lower dimensional latent space. Because genes within a cell have no canonical ordering, Geneformer represents each cell as a rank-value encoding of its genes sorted by expression, emphasizing features that distinguish cellular state while deprioritizing broadly expressed housekeeping genes. This representation strategy is particularly well suited to perturbation modeling: up- or down-regulating a gene corresponds to shifting its position in the rank ordering, and gene deletion can be approximated by removing the corresponding token. In this work, we use token removal as an in silico deletion operator and do not model up- or down-regulation via rank shifts, which would require choosing an arbitrary perturbation magnitude that may not be comparable across heterogeneous cell-line contexts.

#### In silico perturbation yields predictive delta embeddings

We represent each DepMap cell line using the pretrained Geneformer-V1-10M model, trained using 10 million parameters and an input size of 2048, applied to its bulk RNA-seq profile. We define a fixed feature gene universe as the union of the SL label gene set (9,856 from SynLethDB; see Datasets) and the 500 DepMap most highly variable genes (HVGs), yielding 10,032 genes. For each cell line, Geneformer constructs the token sequence by ranking genes by expression and selecting the top-ranked genes up to the context length.

Let **X**_*c*_ ∈ ℝ*d* denote the unperturbed fixed-dimensional *cell-level* embedding. To simulate a knockout of gene *g* in cell line *c*, we remove the token corresponding to *g* from the input sequence and re-embed to obtain a perturbed cell embedding 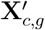. If *g* is not present in the token sequence, we do not generate a perturbed embedding for (*c, g*). We then define the delta embedding

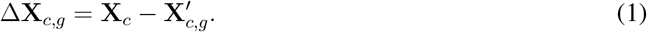

We emphasize that this operation does not simulate a mechanistic biological knockout; rather, ≤**X**_*c,g*_ is a model-defined counterfactual representation shift induced by removing gene *g* from the cell context and re-embedding the remaining transcriptomic context with pretrained Geneformer. Both **X**_*c*_ and 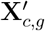 are extracted from the penultimate layer to avoid reliance on the task-specific final layer.

Empirically, ≤**X**_*c,g*_ embeddings capture perturbation-relevant shifts that are predictive of CRISPR viability effects (**Fig**. 2A), and yield features that transfer to SL prediction (**Fig**. 2B, 2C), providing a model-driven proxy for perturbational signaling without relying on curated knowledge graphs. We generate perturbed embeddings (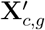) for each cell line and gene combination, for all HVGs and relevant SL and non-SL genes, restricted to genes present in each cell line’s 2048-token sequence. Afterwards, we compute the delta embeddings as the primary input features for Cilantro-sl.

**Figure 1.**
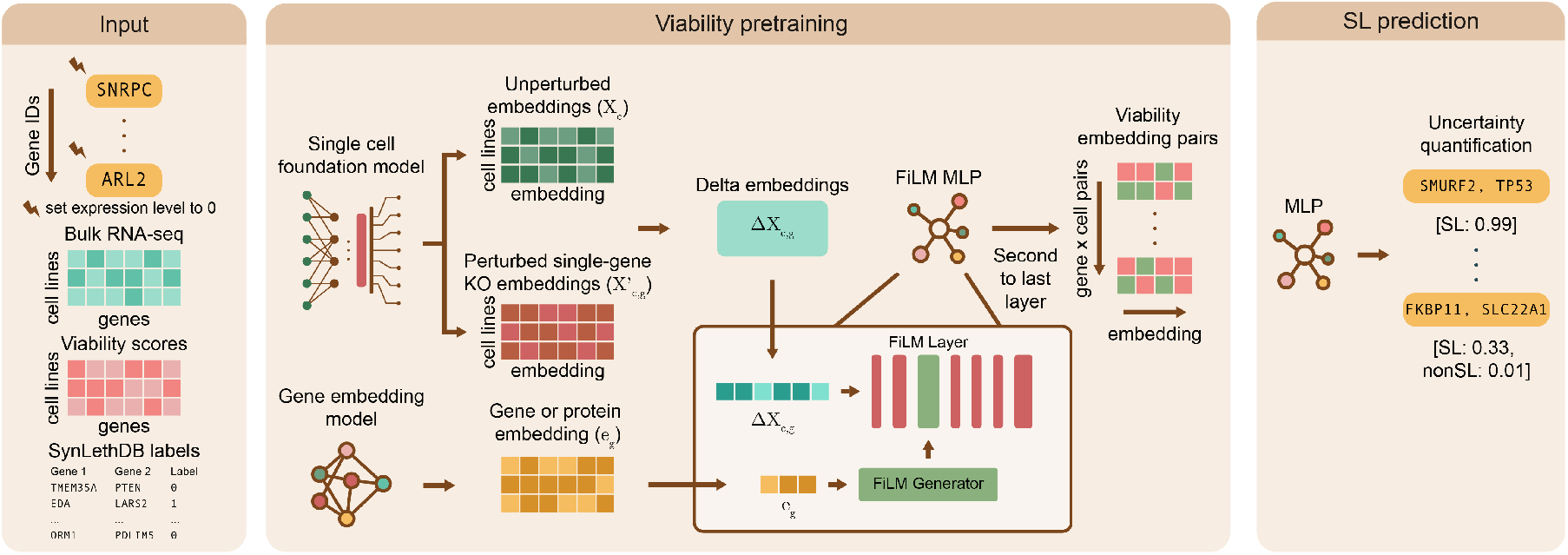
Overview of Cilantro-sl. *Viability pretraining:* Cilantro-sl uses a single-cell foundation model (scFM) to embed each cell line with and without gene knockouts (KOs) and forms delta embeddings by subtracting KO from control embeddings. Gene embeddings for the KO are fused with the delta embeddings using a FiLM layer. Regression pretraining is conducted on DepMap CRISPR data to learn viability embeddings for each gene-cell KO combination. *SL prediction:* For each candidate gene pair, Cilantro-sl combines the corresponding viability embeddings to predict synthetic lethality and applies conformal prediction to obtain calibrated confidence scores. This yields a ranked list of candidate SL pairs with per-pair uncertainty estimates.

**Figure 2.**
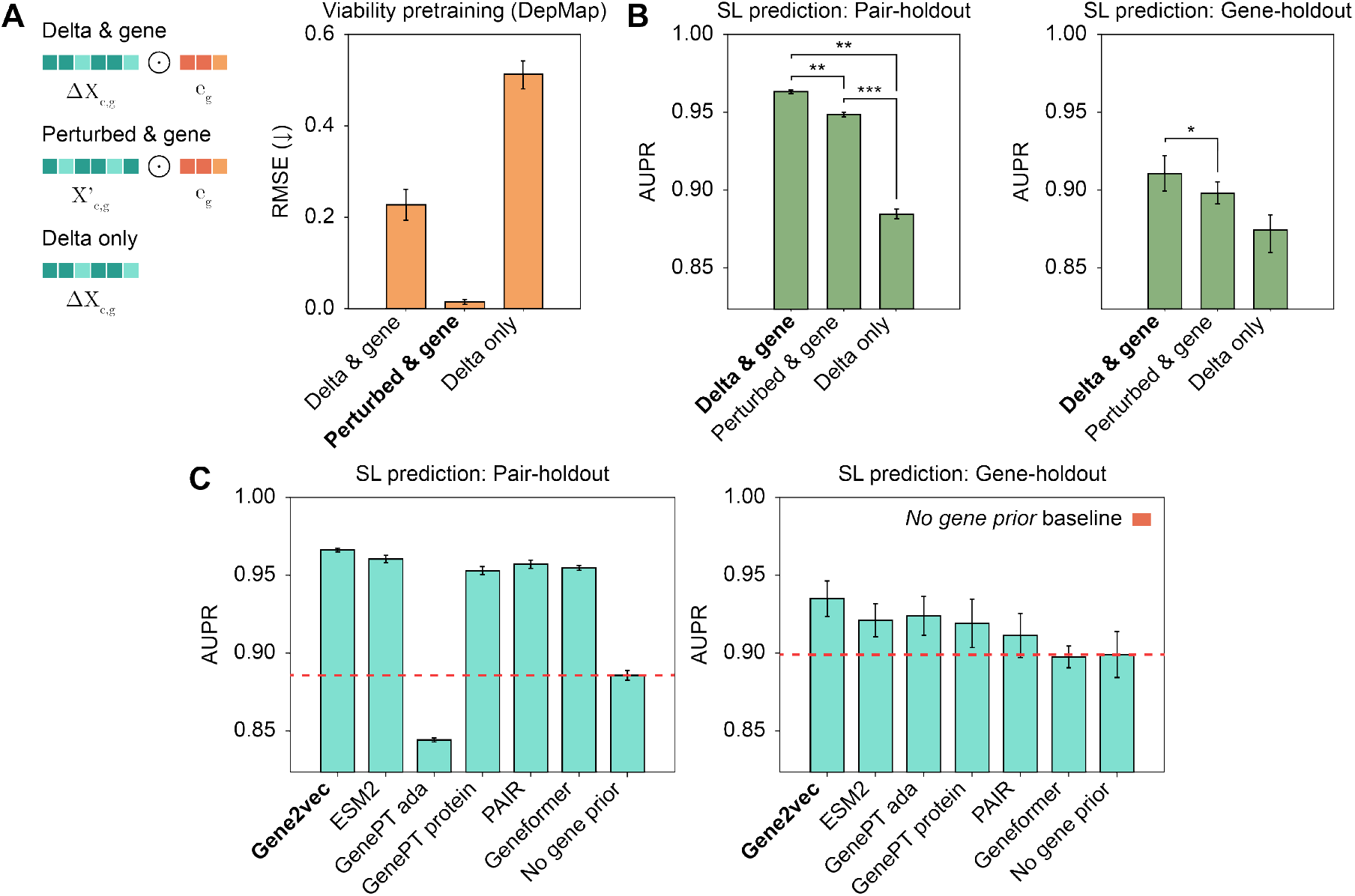
Performance of different inputs to the pretraining stage of Cilantro-sl. **A.** We consider delta embeddings conditioned on Gene2vec (*delta & gene*), perturbed embeddings conditioned on Gene2vec (*perturbed & gene*), and delta embeddings without gene priors (*delta only*). Root mean square error (RMSE) ↓ measures the viability score regression task, with standard deviation computed across different cell lines from the DepMap dataset. **B**. Five-fold cross-validated performance (AUPR) of the three input representations on the Pair-holdout and Gene-holdout SL prediction tasks. Significance was assessed using paired tests (*** : *p* ≤ 0.001, ** : *p* ≤ 0.01, * : *p* ≤ 0.05). **C**. Gene2vec embeddings enhance Cilantro-sl’s predictive capabilities. All gene priors are integrated with Geneformer delta embeddings with FiLM during pretraining. Gene2vec, ESM2, and Geneformer have dimensionality 128 while GenePT ada, GenePT protein, and PAIR have dimensionality 256, all reduced using PCA. *No gene prior* refers to using no gene embedding, i.e. delta only.

##### Gene identity priors via FiLM improves regression

The Geneformer delta embeddings ≤**X**_*c,g*_ provide cell- and context-specific representations of how each KO perturbs a given lineage. However, the delta embeddings do not explicitly encode global information about a gene’s behavior across many contexts. To complement perturbation-aware signal, we introduce a separate gene identity prior **e**_*g*_ learned from large-scale gene co-expression data via Gene2vec embeddings [16]. Here, 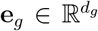 denotes the pretrained Gene2vec embedding for gene g (with *d*_*g*_ = 128), which is kept frozen during training.

Conceptually, this prior serves a similar role to a gene network, but encoded as data-driven pretrained embeddings rather than an explicit, curated graph. Whereas traditional SL methods operate directly on symbolic priors such as PPI networks, GO graphs, or SL knowledge graphs, Cilantro-sl instead relies on learned priors, with Geneformer providing perturbation-sensitive cell embeddings and Gene2vec supplying a global gene-identity representation.

To inject gene identity into the KO representation, we apply a Feature-wise Linear Modulation (FiLM) [18] conditioning layer that modulates the delta embedding by the gene prior. Concretely, for each KO of gene *g* in cell line *c*, we compute scaling and shifting parameters

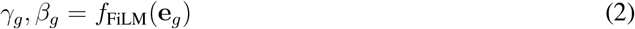

and obtain a FiLM-conditioned representation

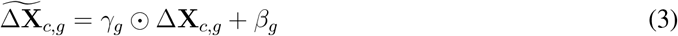

where *f*_FiLM_ is the generator and ⊙ denotes elementwise multiplication. This mid-to-late fusion strategy preserves the structure of the Geneformer delta embeddings while allowing the model to selectively modulate them using the global gene identity information. In this way, Cilantro-sl leverages biological prior knowledge in a learnable manner through the use of gene priors that can be up- or down-weighted by the regression model. Our ablations show that adding Gene2vec priors via FiLM consistently improves KO viability prediction over using delta embeddings alone (**Fig**. S3, **Table** S3).

##### Viability embeddings and SL prediction

We train a lightweight multi-layer perceptron (MLP) on FiLM-modulated features to regress gene-effect viability score, where more negative scores reflect stronger dependency on the gene, using mean-squared error (MSE) loss. We define the viability embedding **V**_*c,g*_ for each (cell line *c*, gene *g*) pair as the 32-dimensional penultimate layer of this network. Because the inputs are bulk RNA-seq measured per cell line, the resulting viability embedding is explicitly cell-line conditioned. Thus, **V**_*c,g*_ summarizes the KO effect in the molecular context of cell line *c*. Within a given cell line, **V**_*c,g*_ is expected to encode the functional impact of gene *g* knockout on cell survival, serving as a compact feature vector for subsequent pairwise SL prediction. This embedding can be interpreted as a transferable perturbation signature that supports zero-shot generalization to unseen genes.

To predict SL interactions, Cilantro-sl trains a second MLP with ReLU activations on the concatenation of viability embeddings [**V**_*c,g*1_; **V**_*c,g*2_] for a given cell line *c* and two KO genes *g*_1_, *g*_2_. Positive SL pairs are obtained from SynLethDB (non-SL pairs are constructed as described in Sec 2.3). Empir-ically, this concatenation retains more signal for SL prediction than the Hadamard product or absolute difference of the viability embeddings (**Table** S4). This formulation enables the model to capture asymmetric or non-additive interactions that cannot be expressed through element-wise operations alone. The MLP is trained for binary SL vs. non-SL classification using weighted cross entropy loss to address label imbalance. By conditioning on the cell line through **V**_*c,g*1_ and **V**_*c,g*2_, Cilantro-sl predicts context-dependent SL interactions rather than global, cell-line-agnostic gene-to-gene relationships. The final layer applies a sigmoid function to output prediction probabilities, which serve as the input for the subsequent conformal prediction module.

### 2.3 Conformal prediction for uncertainty-aware SL selection

SL labels are inherently noisy, as resources such as SynLethDB [19] aggregate evidence from CRISPR screens, RNAi screens, text mining, and computational prediction, leading to variable label quality.

To quantify predictive uncertainty before proposing SL or non-SL pairs for downstream validation, Cilantro-sl applies conformal prediction on top of the SL classifier.

Prediction uncertainty can arise from many sources, including model hyperparameters, label noise, and distributional variation in the data [20]. Conformal prediction addresses this by producing per-sample *prediction sets* with finite-sample coverage guarantees. Under the standard exchangeability assumption that calibration and test points are independent and identically distributed (i.i.d.) or exchangeable, and given a miscoverage rate parameter *α* ∈ (0, 1), conformal prediction ensures that the prediction set contains the true label with probability at least 1 − *α* without requiring parametric assumptions about the model or the underlying data distribution [20–22].

Let *f* (*x*) ∈ ℝ^2^ denote model scores for labels *y* ∈ {0, 1} (non-SL, SL), with softmax probabilities *σ*(*f* (*x*))_*y*_. We define the non-conformity score for input *x* and label *y*:

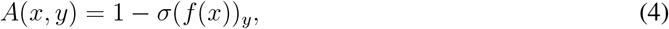

so larger values indicate lower compatibility with the label [22].

On a held-out calibration set 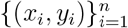, we compute *A*_*i*_ = *A*(*x*_*i*_, *y*_*i*_) for the true label *y*_*i*_, sort the values so that *A*_1_ ≤ … ≤ *A*_*n*_, and for a chosen miscoverage rate *α* define the empirical quantile

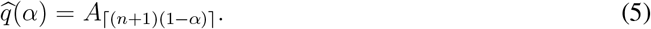

For a test point *x*, we define the prediction set as:

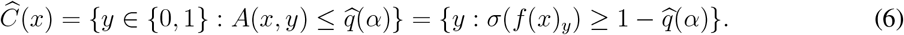

Prediction sets contain a label if its non-conformity score does not exceed 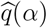, and their size lies in {1, 2} for this binary task. Singleton sets (|*Ĉ* (*x*)| = 1) indicate low ambiguity and high model confidence, whereas larger sets (|*Ĉ* (*x*)| = 2) indicate that the model cannot confidently distinguish SL from non-SL gene pairs. As *α* decreases, desired coverage increases, and prediction sets become more conservative on average.

We also compute conformal *p*-values for each label:

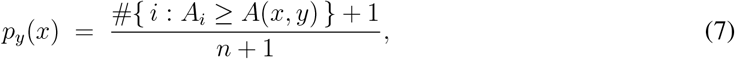

using the standard +1 smoothing. A label is included in the prediction set if and only if *p*_*y*_(*x*) *> α*, which yields the equivalent rule: *Ĉ* (*x*) = {*y* : *p*_*y*_(*x*) *> α*}.

For experimental prioritization, singleton prediction sets at low *α* correspond to high-confidence SL calls. We assign a single *confidence score* to each test sample by taking the maximum conformal *p*-value across both labels: max_*y*_ *p*_*y*_(*x*). High conformal *p*-value (or equivalently, low rank among the calibration scores) indicate high-confidence predictions.

## 3 Results

### 3.1 Datasets

SynLethDB 2.0 [19] contains 35,943 human SL pairs and 2,899 non-SL pairs spanning 9,856 unique genes. Of these, 9,748 are present in both Geneformer and the Cancer Dependency Map (DepMap [23]) and serve as valid inputs to Cilantro-sl. DepMap provides bulk RNA-seq profiles for 1,408 cancer cell lines across 19,194 protein-coding genes, and CRISPR gene-effect viability scores for 1,078 cell lines and 17,453 genes, where more negative values indicate stronger dependency. Intersecting RNA-seq and viability data yields 1,021 overlapping cell lines. We define the feature gene set as the union of the 9,856 SynLethDB SL/non-SL genes (restricted to the 9,748 which are present in DepMap) and the 500 most highly variable genes determined from DepMap RNA-seq for a total of 10,032 unique genes.

To expand the limited set of annotated non-SL pairs, we constructed a correlation-aware negative sampling set using DepMap expression data, prioritizing low-correlation gene pairs as described previously [24]. Negatives were restricted to pairs within the defined feature set and sampled under both 1× and 5× negative-to-positive ratios. The resulting datasets contain 22,081 negatives in the 1× setting and 130,378 negatives in the 5× setting. After viability pretraining, the downstream SL classification corpus comprises 6,101,812 cell-pair samples in the 1× setting and 11,145,517 cell-pair samples in the 5× setting. SynLethDB provides incomplete and inconsistently matched cell-line annotations, with only 13 DepMap cell lines shared between positive and negative labels. Because CILANTRO-SL generates cell-line–specific predictions, we therefore evaluate at the gene-pair level by averaging the cell-line–specific scores into a single pair-level prediction for benchmarking. We primarily evaluate Cilantro-sl and all benchmarks on the 5× setting, as SL interactions are sparse relative to the combinatorial space of gene pairs, leading to a strongly negative-dominant distribution in practical screening contexts.

#### Data split methods

We consider two cross-validation regimes:

- **Pair-holdout (seen-gene, unseen-pair)**. No gene pair (*g*_*i*_, *g*_*j*_) appears in both train and test, while individual genes may appear in both. This assesses generalization to novel combinations among previously seen genes.
- **Gene-holdout (unseen-gene)**. Genes are partitioned into distinct sets *G*_train_ and *G*_test_. Test pairs satisfy *g*_*i*_, *g*_*j*_ ∈ *G*_test_. This is a zero-shot evaluation on novel genes and novel relationships and simulates the regime where curated PPI and GO resources are especially sparse.

### 3.2 Optimizing viability-pretrained representations for SL prediction

To isolate which representation choices most strongly support SL prediction, we conducted a controlled ablation over three factors: (i) the Geneformer representation used for viability pretraining (perturbed 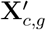 versus delta Δ**X**_*c,g*_), (ii) the presence or absence of a gene identity prior, and (iii) the fusion strategy used to condition on that prior. Unless otherwise stated, the gene prior is the Gene2vec co-expression embedding reduced to 128-dimensions with PCA. Gene2vec with 128-dimensional and 64-dimensional reductions performed comparably and outperformed 32-dimensional reductions (**Table** S1). As a no-pretraining control, we projected Δ**X**_*c,g*_ to 32-dimensions via PCA to match the size of the pretrained viability embedding.

On the viability regression pretraining task, conditioning on a gene prior yields the largest gains (**Fig**. 2A). Delta embeddings conditioned on Gene2vec achieve a root mean square error (RMSE) of 0.230 *±* 0.000, a Spearman correlation of 0.864 *±* 0.001 and *R*^2^ of 0.85 *±* 0.002 (**Fig**. S1). The perturbed embedding 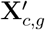 conditioned on a gene prior attain near-perfect fit in all three metrics with negligible variance. By contrast, removing the prior (*delta only*) results in poor fit (*R*^2^ = −0.255 *±* 0.020), indicating that unconditioned Geneformer embeddings alone are insufficient to capture gene-specific knockout effects.

When transferred to SL pair classification, however, the ordering reverses: viability embeddings trained with Δ**X**_*c,g*_ generalize more effectively than those trained with 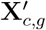 (**Fig**. 2B). Relative to 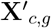, Δ**X**_*c,g*_ improves Pair-holdout by +1.62% AUPR, +6.24% AUC, and +3.11% F1, and improves Gene-holdout by +1.26% AUPR, +5.71% AUC, and +5.24% F1 (**Fig**. S2). Paired t-tests between the delta & gene variant and the perturbed & gene yield *t* = 6.534, *p* = 0.003 in Pair-holdout and *t* = 3.481, *p* = 0.025 in Gene-holdout, indicating significant improvements. In both regimes, the delta-only variant is uniformly worst. Together, these results suggest that while *perturbed & gene* best fits the pretraining regression objective, it overfits viability, whereas *delta & gene* preserves features that are more transferable to SL predictions.

We next examined how the gene prior should be integrated into the model. Mid-to-late stage fusion with FiLM modestly but consistently outperforms early fusion by concatenation for both Gene2vec and ESM2 priors, yielding modest AUPR gains and larger F1 improvements in both Pair- and Gene-holdout (**Table** S3).

Finally, we compared multiple gene-identity priors for conditioning Geneformer delta embeddings during viability pretraining. We compared (i) no prior, (ii) Gene2vec [16], (iii) ESM2 [14], (iv) GenePT variants – text-derived gene embeddings generated from GPT (GenePT ada; GenePT protein) [15], (v) PAIR – protein language model embeddings (ESM2-initialized variant) finetuned with associated text annotations to couple sequence/structure and textual modalities [25], and (vi) Geneformer’s own gene embeddings. Integrating Gene2vec yields the highest AUPR among all priors, with a +0.58% improvement over ESM2 in Pair-holdout and +1.39% in Gene-holdout (**Fig**. 2C). Removing gene information substantially degrades SL prediction, with AUPR decreasing by more than 3.75% in Pair-holdout. Across both zero-shot regimes, Cilantro-sl performs best with Gene2vec or ESM2, outperforming text-based GenePT embeddings, PAIR, and Geneformer’s own gene embeddings in F1, precision, and recall (**Fig**. S3) across both prediction tasks, although all priors improve over the no-gene prior baseline.

### 3.3 Benchmarking Cilantro-sl for SL prediction

We benchmarked Cilantro-sl against established SL prediction frameworks spanning graph-based, matrix factorization, and pretrained representation paradigms. Graph-based models included KG4SL [6], SLMGAE [8], DDGCN [5], and MVGCN-iSL [7], which leverage curated knowledge graphs, PPI, or GO similarity networks. SL^2^MF [9] represents matrix factorization approaches incorporating such priors, and ESM4SL [11] is a pretrained protein-embedding–based method. To disentangle representational gains from classifier capacity, we additionally evaluated an MLP variant of ESM4SL, which we call ESM+MLP. All models were trained and evaluated on the same SynLethDB-derived dataset and sampling configuration (Section 3.1) under Pair-holdout and Gene-holdout zero-shot splits. Given the pronounced label imbalance inherent to SL prediction, we report five-fold cross-validated area under the precision-recall curve (AUPR) together with F1 (**Fig**. S4).

Cilantro-sl achieves the strongest overall performance in the more stringent zero-shot Gene-holdout setting, where each gene in the test set is unseen during training. In this regime, Cilantro-sl achieves the highest F1 score among evaluated methods while maintaining competitive AUPR (0.7148 versus 0.7325 for KG4SL) (**Fig**. 3A). Relative to KG4SL and ESM4SL, Cilantro-sl improves Gene-holdout F1 by 28.6% and 49.9%, respectively. DDGCN cannot be evaluated in this setting, as it derives node features directly from the SL adjacency and therefore cannot construct representations for completely unseen genes. To assess robustness to gene-level distribution shift, we compared performance between Pair-holdout and Gene-holdout using the average (AUPR + F1)*/*2 (**Fig**. 3B). Although ESM4SL achieves the strongest performance under Pair-holdout, its F1 decreases from 0.914 to 0.453 under Gene-holdout, whereas Cilantro-sl exhibits a smaller reduction (0.18 versus 0.46 absolute decrease), indicating improved stability under gene-level distribution shift. Across metrics, Cilantro-sl shows the most balanced aggregate performance in the Gene-holdout setting (**Fig**. S4B), whereas several baselines exhibit marked degradation.

**Figure 3.**
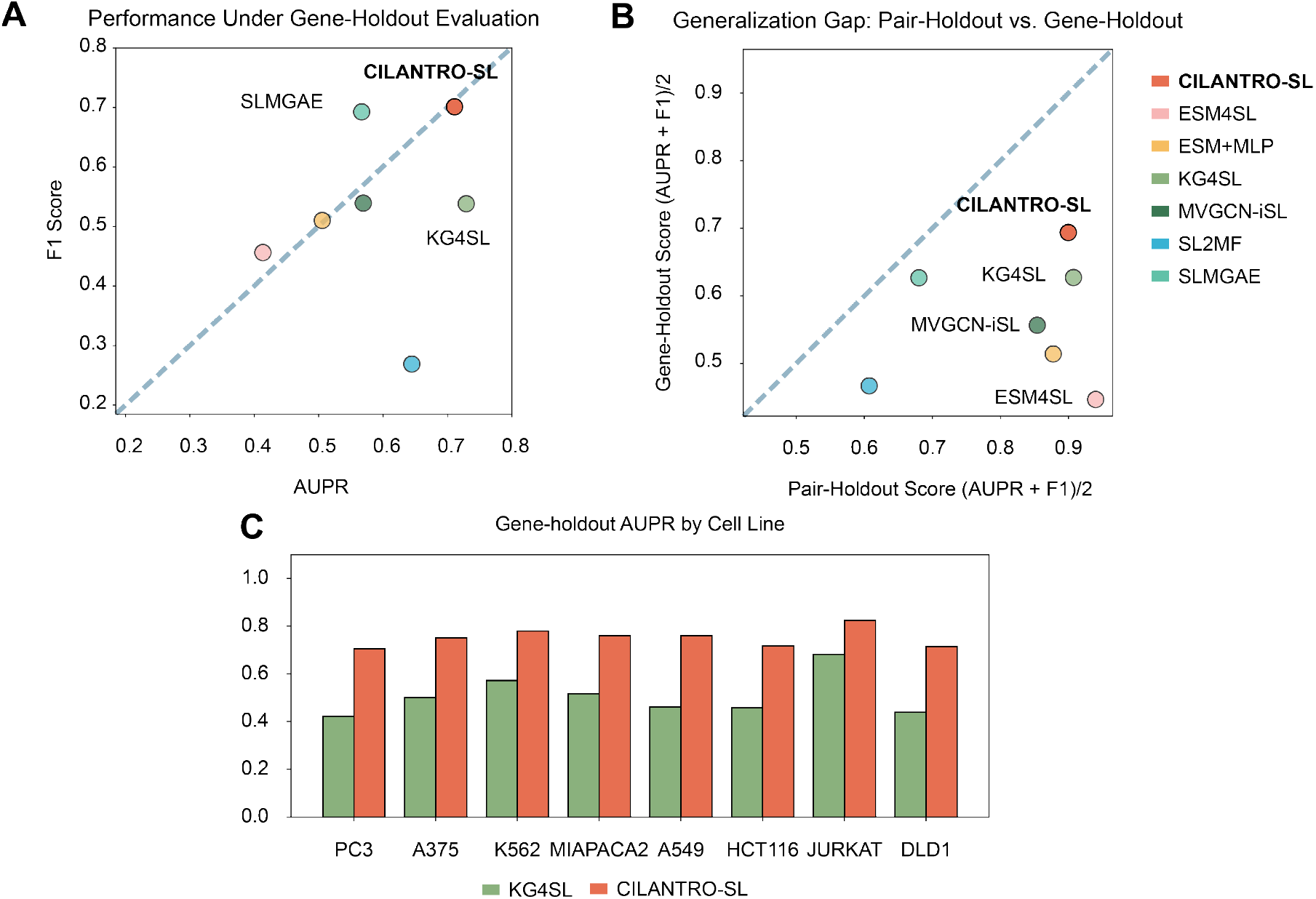
Five-fold cross-validated performance evaluation (AUPR and F1 score) of Cilantro-sl against baseline methods KG4SL, SL^2^MF, SLMGAE, DDGCN, MVGCN-iSL, ESM4SL, and ESM+MLP. **A.** F1 Score vs. AUPR on the harder Gene-holdout task under 5 × negative sampling. **B**. Average score ((AUPR + F1) / 2) comparison of Gene-holdout against Pair-holdout under 5 × negative sampling. Further deviation below the line *y* = *x* implies that the model performs much better on the easier Pair-holdout task than the Gene-holdout one. **C**. Performance comparison for cell-line-specific prediction for KG4SL and Cilantro-sl under the Gene-holdout split strategy.

Given the context-dependent nature of synthetic lethality, we next conducted a cell-line-stratified Gene-holdout analysis across eight well-sampled cell lines (A375, A549, DLD1, JURKAT, K562, HCT116, PC3, and MIAPACA2). Cilantro-sl exceeds KG4SL in AUPR within each cell line (**Fig**. 3C), demonstrating that the observed performance gains from Cilantro-sl generalize across cellular backgrounds. Consistent with our primary 5× results, Cilantro-sl exhibits comparatively smaller degradation when moving from 1× to the more challenging 5× negative sampling regime across both AUPR and F1 relative to several baselines (**Fig**. S7A-B). Together, these findings indicate that viability-pretrained, foundation-model-derived representations provide an effective alternative to static interaction topology and pre-trained protein embeddings, enabling stable SL prediction that generalizes to previously unseen genes while remaining robust across cellular contexts.

### 3.4 Conformal prediction-based uncertainty quantification

To assess the reliability of SL predictions, we applied conformal prediction on top of Cilantro-sl and evaluated coverage, set size, and confidence stratification across five cross-validation folds. Rather than producing fixed-threshold scores, conformal prediction enables user-controlled reliability: by specifying a target error level *α*, users obtain prediction sets that achieve marginal coverage of at least 1 − *α*. In each fold, Cilantro-sl was trained on the training split, calibrated conformal thresholds on a held-out 10% subset, and evaluated performance on an independent test set. Empirical coverage closely matched the target coverage level 1 − *α* across a broad range of *α* values (**Fig**. 4B). The observed coverage followed the *y* = *x* diagonal, indicating that prediction sets achieve the intended finite-sample error control rather than being systematically conservative or under-confident. These results demonstrate that CILANTRO-SL produces well-calibrated prediction sets in practice, satisfying marginal coverage guarantees under standard assumptions [26]

**Figure 4.**
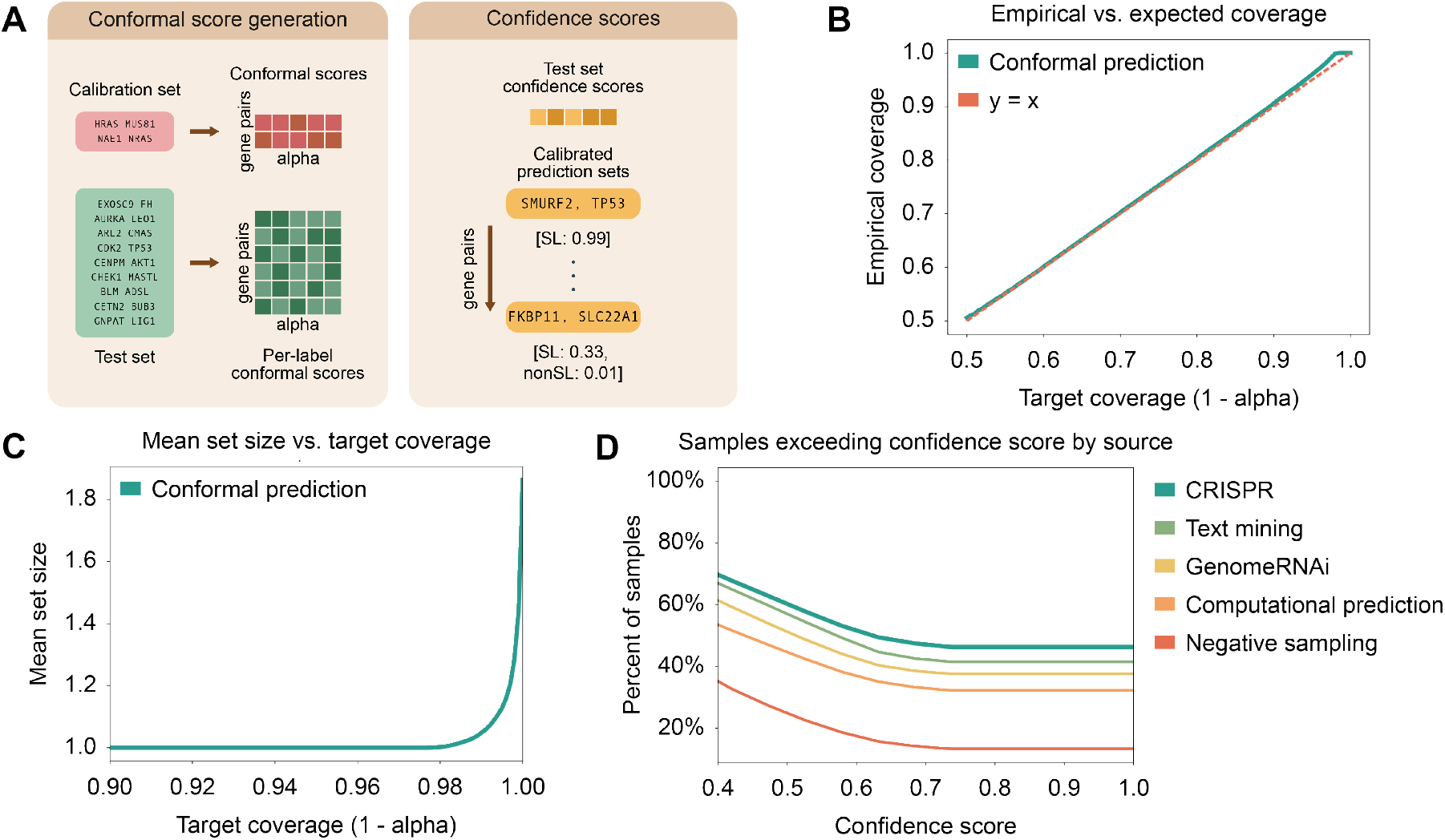
Evaluation of the uncertainty quantification stage of Cilantro-sl under Pair-holdout. **A.** Overview of conformal prediction confidence score and prediction set generation process. **B**. Empirical coverage of true labels in prediction sets compared to the target coverage (1 − *α*). **C**. Measurement of the mean prediction set size against target coverage. **D**. Stratification of prediction confidence based on SL label source. We report the percentage of samples coming from a certain source (e.g. CRISPR) that has a confidence score exceeding a certain threshold.

We next examined the tradeoff between coverage and prediction set size. Increasing the target coverage resulted in larger prediction sets, as achieving a lower tolerated error rate (smaller *α*) requires relaxing the non-conformity threshold, to include additional candidate labels (**Fig**. 4C). Notably, CILANTRO-SL maintains prediction sets of size one for nearly all samples up to a target coverage of roughly 98%, indicating that the model remains highly decisive even under stringent coverage guarantees.

Because SL labels are heterogeneous, we further evaluated Cilantro-sl’s confidence as a function of the annotation source in SynLethDB, which aggregates gene pairs supported by CRISPR knock-out screens, previous computational predictions, literature-derived evidence, and genome-wide RNAi screens. CRISPR-supported pairs exhibited the highest proportion of top-confidence predictions, with 46.28% achieving the highest confidence score (**Fig**. 4D). Text-mined pairs also showed substantial enrichment in high-confidence bins (41.50%), consistent with partial grounding in prior experimental observations reported in the literature. Crucially, Cilantro-sl’s Stage 1 pretraining leverages single-gene CRISPR viability measurements but does not incorporate pairwise SL annotations. Therefore, the strong enrichment of CRISPR-supported pairs in high-confidence bins suggests that our perturbation-informed representations successfully capture functional dependencies that transfer to pairwise synthetic lethality prediction. Across confidence bins, the proportion of SL relative to non-SL pairs increases with confidence, indicating that conformal scores meaningfully stratify prediction reliability (Fig. S5). CRISPR-supported pairs are enriched in the highest-confidence bins relative to lower-confidence bins, whereas computationally inferred labels exhibit the opposite trend. In contrast, negatively sampled pairs constructed without direct experimental support receive much lower confidence: only 13.34% of negatively sampled pairs achieve the highest confidence score, and more than half receive confidence below 40%. Together, these findings demonstrate that Cilantro-sl provides calibrated uncertainty estimates grounded in the perturbation-informed embeddings learned during viability pretraining. These representations enable reliable confidence estimation and principled prioritization of candidate SL interactions for downstream validation.

### 3.5 Biological enrichment and prioritization of high-confidence SL predictions

We next asked whether high-confidence SL predictions cluster in coherent pathways and whether this varies by evidence source (Supplementary Section G). Among genes participating in high-confidence CRISPR-anchored pairs (493 genes; conformal confidence ≥ 0.9), GO enrichment revealed strong signal for mitochondrial translation, translational control, cell-cycle progression, and checkpoint regulation (**Fig**. S6A). Top terms included mitochondrial translational elongation and termination, cellular macro-molecule biosynthesis, protein phosphorylation, regulation of p53 signaling, DNA metabolic process, and mitotic cell-cycle phase transition (all FDR ≪ 10^−10^).

Genes appearing only in high-confidence computational-prediction pairs (1,379 genes) showed even sharper enrichment for DNA damage response and mitotic control (**Fig**. S6B). Highly significant terms included DNA metabolic process, DNA repair, mRNA processing and spliceosome function, cellular response to DNA damage stimulus, DNA replication, mitotic spindle organization, G1/S transition of the mitotic cell cycle, and double-strand break repair (FDR down to *<* 10^−50^). Thus, both strata concentrate in cancer-relevant pathways, with computational-only predictions preferentially highlighting DNA repair, replication, and mitotic programs that are natural candidates for novel SL interactions.

To move from pathway-level trends to concrete hypotheses, we focused on computational-only high-confidence pairs whose partners shared at least one DNA-damage response, replication, mitotic, or RNA-splicing term and ranked them by supported cell-line contexts and conformal confidence.

Among the top-ranked candidates, several gene pairs stood out for their cancer relevance and therapeutic tractability. Notably, TP53-PARP1 links the most frequently mutated tumor suppressor in human cancers [27] to a DNA repair enzyme targeted by FDA-approved *PARP1*-inhibitors [28, 29]. Loss of *TP53* disables the G1 checkpoint and increases reliance on S/G2-phase DNA damage response (DDR) programs, while *PARP1* plays a key role in repairing replication-associated DNA lesions, making TP53-PARP1 a mechanistically plausible DDR-focused candidate. This illustrates how our high-confidence set on SL gene pairs from SynLethDB can recover biologically coherent DDR interactions whose components lie on well-studied PARP-inhibitor pathways but are not CRISPR-validated, and therefore hold promise for future experimental investigation.

As a complementary example, AURKA-BUB1B links a mitotic kinase that promotes centrosome maturation and spindle assembly to a core spindle assembly checkpoint protein that prevents chromosome missegregation [30, 31]. This pair is annotated with shared GO terms related to mitotic spindle organization and mitotic cell-cycle phase transitions, and *AURKA* is frequently overexpressed in human cancers and has small-molecule inhibitors in preclinical and clinical testing [30]. Although AURKA-BUB1B has no CRISPR-based validation in SynLethDB, its convergence on mitotic spindle and check-point, together with the cancer relevance and clinical tractability of *AURKA*, makes it a biologically compelling candidate for future experimental follow-up. Together, these examples illustrate how our high-confidence set of SL gene pairs derived from SynLethDB can recover mechanistically coherent interactions involving well-studied cancer genes and drug targets beyond those supported by CRISPR evidence.

## 4 Discussion

We introduced Cilantro-sl, an uncertainty-aware framework for SL prediction that replaces reliance on static interaction graph priors with pretrained, context-conditioned biological representations. The model learns single-gene viability embeddings that integrate in silico Geneformer perturbation shifts with Gene2vec priors, capturing cell-line-specific functional signals without dependence on predefined interaction topology. Conformal prediction further provides calibrated, per-pair confidence estimates to support reliable downstream prioritization.

Across both Pair-holdout and the stricter Gene-holdout setting, Cilantro-sl demonstrates strong performance relative to graph-based and pretrained representation baselines. Importantly, performance degradation from Pair-holdout to Gene-holdout is comparatively small, indicating robustness when predicting interactions involving previously unseen genes. Cell-line–stratified analyses further show consistent gains across diverse cancer contexts, supporting the model’s context-conditioned design.

The modular design of Cilantro-sl naturally supports incorporating multiple pretrained priors in parallel, beyond Gene2vec and Geneformer. For example, protein language model embeddings and text-derived gene representations can be incorporated as additional gene-level priors to further strengthen the viability embeddings. Moreover, our single KO-based viability formulation can be extended to modeling higher-order perturbations beyond SL (e.g., combinatorial KOs) [32], enabling Cilantro-sl to exploit multiple simultaneous priors even in more complex perturbation settings.

Cilantro-sl is well suited as a screening tool for precision oncology. By generating cell-line conditioned scores and ranking gene pairs with calibrated confidence, the framework supports prioritization of high-value candidates for experimental validation. Explicit conditioning on cellular context also enables more targeted and interpretable hypothesis generation in specific tissues or molecular backgrounds. By combining pretrained biological foundation models with uncertainty quantification, CILANTRO-SL directly addresses core limitations of data scarcity and reliance on static curated features that constrain current SL methods. In doing so, Cilantro-sl illustrates how future SL predictors intended for experimental screening should integrate uncertainty by design to deliver more confident, actionable SL hypotheses for experimental and translational follow-up.

## Acknowledgment

This work was supported, in part, by National Institutes of Health Common Fund 4D Nucleome Program grant UM1HG011593 (J.M.); National Institutes of Health Common Fund Cellular Senescence Network Program grant UH3CA268202 (J.M.); and National Institutes of Health grants R01HG007352 (J.M.), R01HG012303 (J.M.), R21DA061481 (J.M.), R03OD039980 (J.M.), and U24HG012070 (J.M.). J.M. was additionally supported by the Ray and Stephanie Lane Professorship, a Guggenheim Fellowship from the John Simon Guggenheim Memorial Foundation, and a Google Research Award. The funders had no role in study design, data collection and analysis, decision to publish or preparation of the manuscript.

## Code Availability

Source code for Cilantro-sl is available at: https://github.com/kaileyhh/Cilantro-SL.

## Author Contributions

Conceptualization, K.H., E.H., J.M.; Methodology, K.H., E.H., J.M.; Software, K.H., E.H.; Investigation, K.H., E.H., J.M.; Writing, K.H., E.H., J.M.; Funding Acquisition, J.M.

