## Supplemental Information for "Uncertainty-aware synthetic lethality prediction with pretrained foundation models"

#### A Choice of gene embedding dimensionality

| | Pair-holdout | | | Gene-holdout | | |
|---|---|---|---|---|---|---|
| **Dimension** | **Precision** | **Recall** | **F1** | **Precision** | **Recall** | **F1** |
| 32 | 0.968 | 0.934 | 0.951 | 0.923 | 0.795 | 0.853 |
| 64 | 0.970 | **0.941** | **0.955** | **0.919** | **0.851** | **0.883** |
| 128 | **0.972** | 0.937 | 0.954 | 0.918 | 0.845 | 0.879 |

**Table S1:** Ablation study on gene embedding dimensionality for Gene2vec, integrated with Geneformer embedding with FiLM during the pretraining stage. All of the dimensionality reductions were computed using PCA.

| | Pair-holdout | | | Gene-holdout | | |
|---|---|---|---|---|---|---|
| **Dimension** | **Precision** | **Recall** | **F1** | **Precision** | **Recall** | **F1** |
| 64 | 0.964 | 0.902 | 0.932 | 0.896 | 0.774 | 0.829 |
| 128 | 0.968 | **0.921** | **0.944** | 0.902 | **0.847** | **0.873** |
| 256 | **0.969** | 0.910 | 0.939 | **0.907** | 0.807 | 0.852 |
| 512 | 0.897 | 0.603 | 0.721 | 0.894 | 0.602 | 0.718 |

**Table S2:** Ablation study on gene embedding dimensionality for ESM2, integrated with Geneformer embedding with FiLM during the pretraining stage. All of the dimensionality reductions were computed using PCA.

#### B Effects of feature-wise linear modulation

| | Pair-holdout | | | Gene-holdout | | |
|---|---|---|---|---|---|---|
| **Fusion method** | **AUC** | **AUPR** | **F1** | **AUC** | **AUPR** | **F1** |
| Gene2vec early fusion | 0.862 | 0.963 | 0.917 | 0.647 | 0.909 | 0.874 |
| **Gene2vec with FiLM** | **0.875** | **0.966** | **0.954** | **0.659** | **0.912** | **0.879** |
| ESM2 early fusion | 0.749 | 0.941 | 0.885 | 0.626 | 0.904 | 0.810 |
| ESM2 with FiLM | 0.856 | 0.960 | 0.939 | 0.599 | 0.898 | 0.852 |

**Table S3:** Feature-wise linear modulation improves five-fold cross-validated performance of CILANTRO-SL. We compare performance with early fusion and mid / late (FiLM) information fusion in the viability pretraining stage across the Pair-holdout and Gene-holdout prediction tasks using both Gene2vec and ESM2 embeddings as the gene prior.

#### C Viability embedding combination methods for CILANTRO-SL

| | Pair-holdout | | | Gene-holdout | | |
|---|---|---|---|---|---|---|
| **Combination method** | **Precision** | **Recall** | **F1** | **Precision** | **Recall** | **F1** |
| Concatenation | **0.972** | **0.937** | **0.954** | **0.918** | **0.845** | **0.879** |
| Hadamard product | 0.945 | 0.847 | 0.894 | 0.911 | 0.795 | 0.849 |
| Absolute difference | 0.933 | 0.822 | 0.874 | 0.906 | 0.737 | 0.812 |

**Table S4:** Ablation study on method of combining viability embeddings to construct the input to the MLP in the SL classification stage of CILANTRO-SL. All viability embeddings were produced with Geneformer delta embeddings conditioned on Gene2vec gene embeddings (dimensionality 128) using FiLM in the pretraining stage of CILANTRO-SL. The concatenation method creates a 64-dimensional input while the hadamard product and absolute difference make 32-dimensional inputs.

#### D Choice of viability pretraining input representations and priors

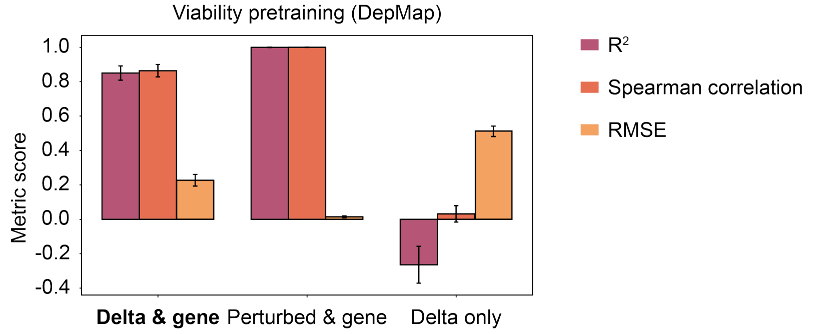


**Figure S1:** Performance (R-squared ↑, Spearman correlation ↑, and root mean square error (RMSE) ↓) of different inputs to the pretraining stage of CILANTRO-SL: delta embeddings conditioned on Gene2vec (*delta & gene*), perturbed embeddings conditioned on Gene2vec (*perturbed & gene*), and delta embeddings without gene priors (*delta only*), with standard deviation computed across different cell lines from the DepMap dataset.

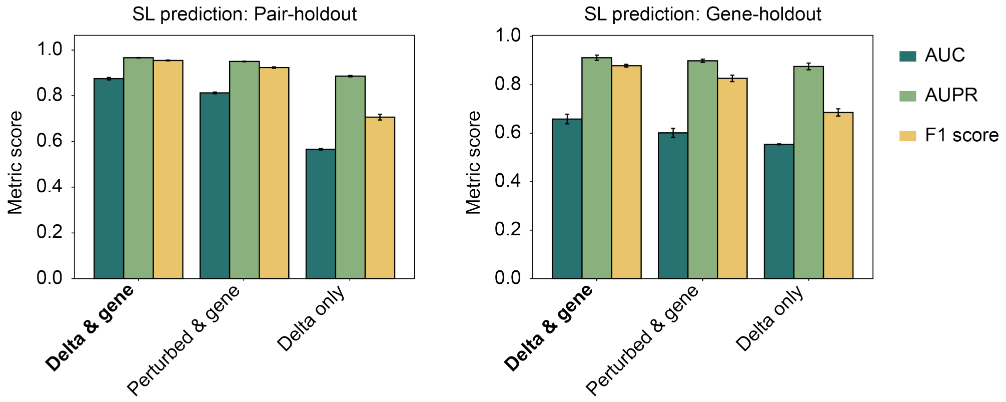


**Figure S2:** Five-fold cross validated performance (AUC, AUPR, F1) of different inputs to the pretraining stage of CILANTRO-SL on the SL classification task: delta embeddings conditioned on Gene2vec (*delta & gene*), perturbed embeddings conditioned on Gene2vec (*perturbed & gene*), and delta embeddings without gene priors (*delta only*).

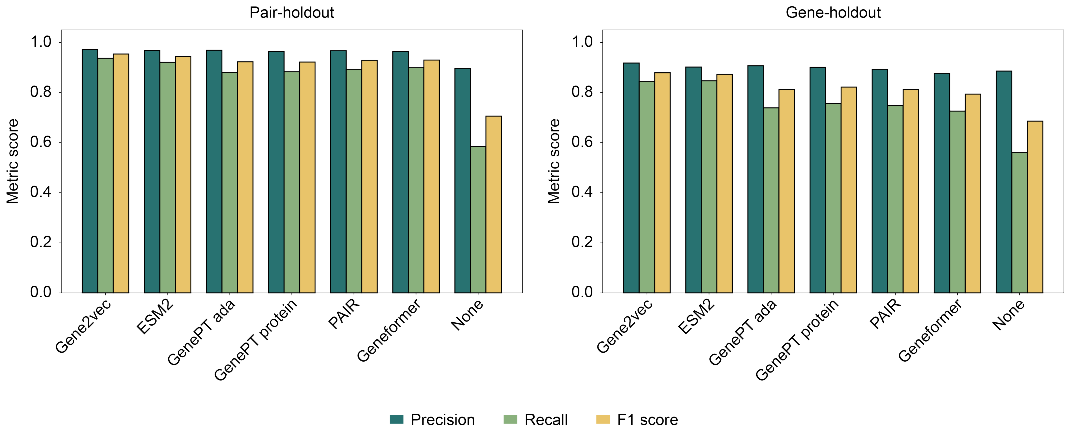


**Figure S3:** Ablation study on choice of gene embedding or protein embedding, integrated with Geneformer delta embeddings with FiLM during pretraining. ESM2, Gene2vec, Geneformer have dimensionality 128, reduced using PCA, while GenePT ada, GenePT protein, and PAIR have dimensions reduced to 256, all using PCA. *None* refers to using no gene embedding, i.e. delta only.

#### E Performance of CILANTRO-SL against baseline methods

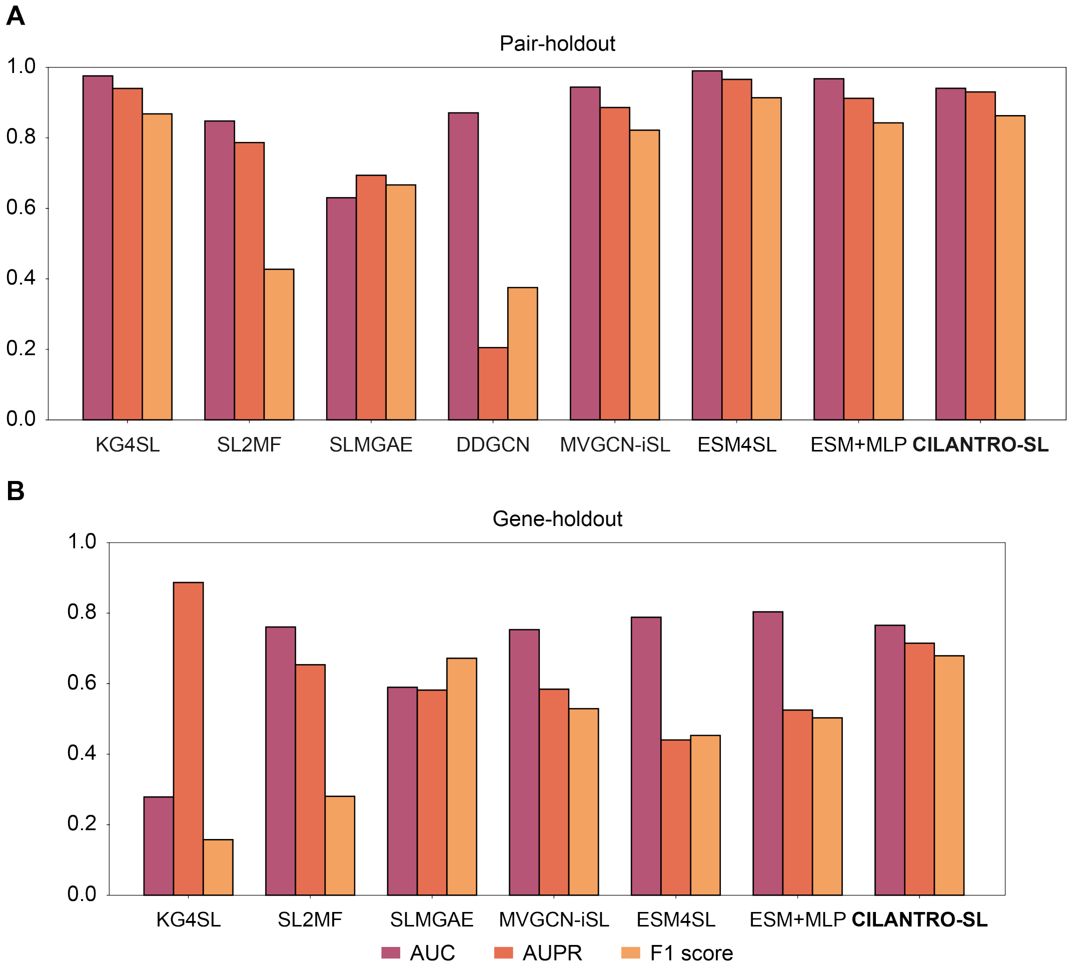


**Figure S4:** Five-fold cross validated performance evaluation (AUC, AUPR, F1) of CILANTRO-SL against baseline methods KG4SL, SL$^2$MF, SLMGAE, DDGCN, MVGCN-ISL, ESM4SL, and ESM+MLP for both data split strategies, Pair-holdout and Gene-holdout.

### F SL pair sample sources by confidence score

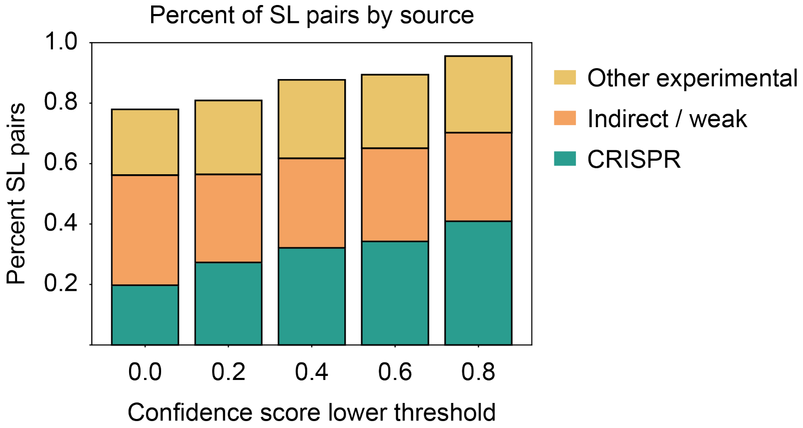


**Figure S5:** Percentage of SL pairs in each confidence score bucket (scores from lower threshold $k$ to upper threshold $k + 0.2$) by SynLethDB sampling source. The *CRISPR* category is gene pairs identified by CRISPR screens, while the *other experimental* category includes GenomeRNAi, RNAi screens, drug screens, high/low throughput sequencing, and Decipher (shRNA screens). The *indirect/weak* category refers to methods such as text mining and computational prediction.

#### G Gene ontology pathway enrichment

We performed Gene Ontology (GO) enrichment to characterize pathways associated with high-confidence SL predictions by evidence source. Starting from the SynLethDB-derived SL pair table with model-based confidence scores, we defined high-confidence pairs as those with conformal confidence $\geq 0.9$ and partitioned them into (i) CRISPR-anchored pairs, where SynLethDB evidence field contained a CRISPR label, and (ii) computational-only pairs, where the evidence contained Computational Prediction but no CRISPR annotation. For each stratum, we constructed a foreground gene set as the union of all genes appearing as either partner in the corresponding high-confidence pairs. GO Biological Process enrichment was then performed using the Enrichr interface implemented in `gseapy` (`gp.enrichr`) with the `GO_Biological_Process_2021` library and Enrichr's default background universe (all genes annotated in the library). Enrichr reports one-sided Fisher's exact test p-values and Benjamini-Hochberg-corrected p-value in the column labeled Adjusted P-value. We considered terms significant if their adjusted p-value was $< 0.05$ and ranked significant terms by this value for visualization. The top enriched terms for each evidence stratum are shown in **Fig.** S6.

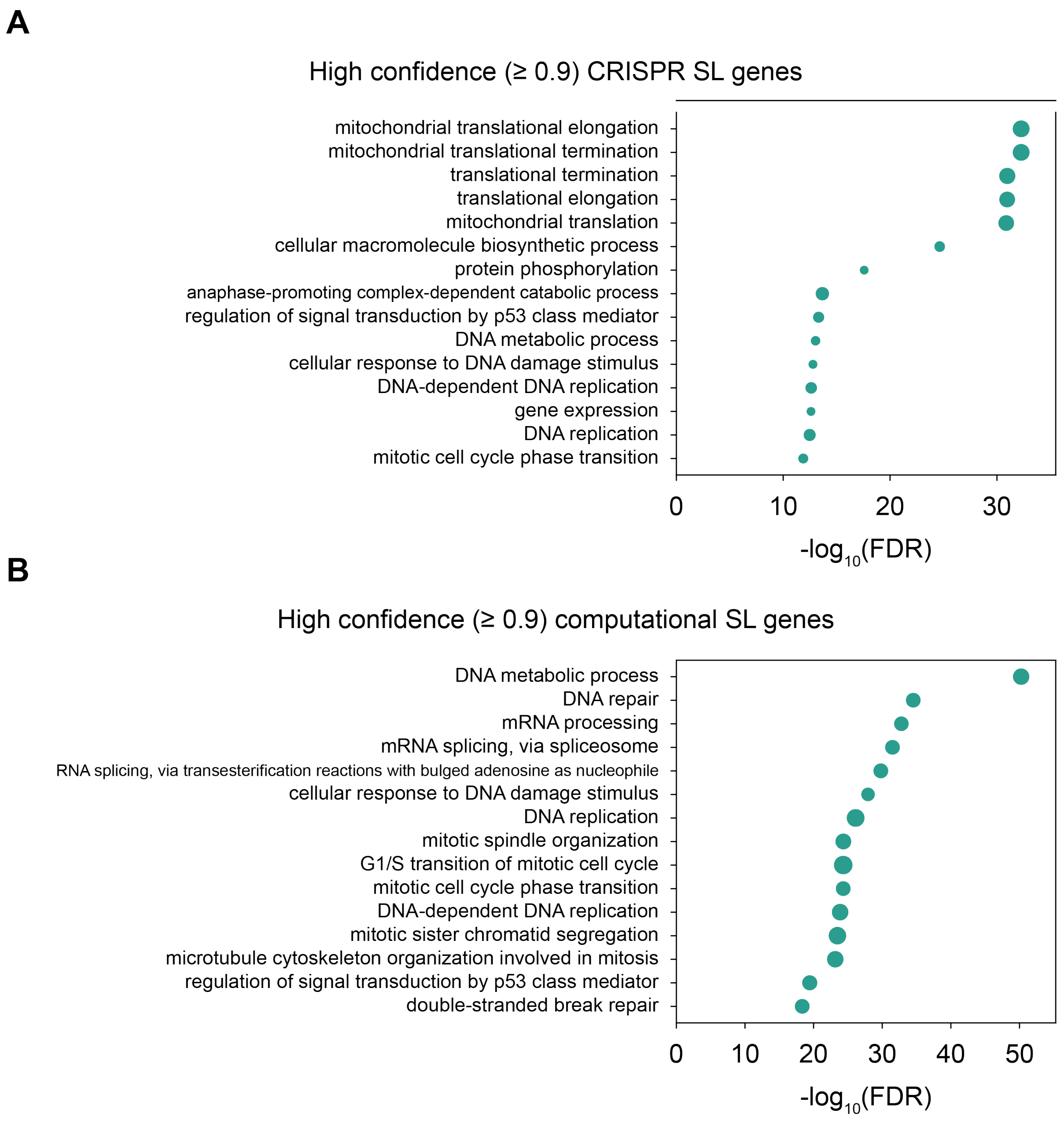


**Figure S6: Gene ontology pathway enrichment data for known SynLethDB SL pairs. A.** Pairs sourced from CRISPR screens where CILANTRO-SL predicts the positive label with a confidence score of at least $0.9$. **B.** Pairs sourced from prior computational prediction methods where CILANTRO-SL predicts the positive label with a confidence score of at least $0.9$.

#### H Robustness to negative sampling by metric

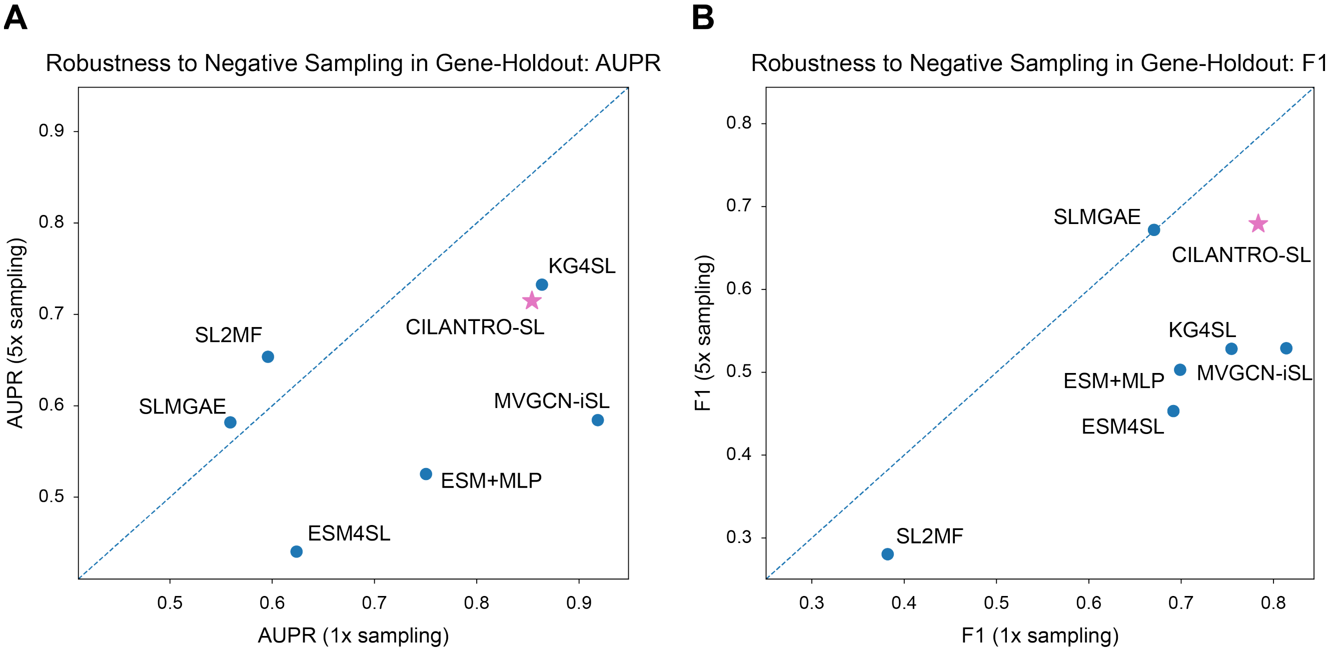


**Figure S7: Robustness to different negative sampling strategies.** Performance of CILANTRO-SL against baseline models of KG4SL, $SL^2MF$, SLMGAE, MVGCN-ISL, ESM4SL, and ESM+MLP, under 1x and 5x negative sampling. Baseline DDGCN was not included as it does not take in negative samples, and has low AUPR and F1 scores (0.20518 and 0.37556 respectively). **A.** AUPR of all methods in the 5x sampling compared to 1x sampling. **B.** F1 of all methods in the 5x sampling compared to 1x sampling.

### I Gene-holdout and pair-holdout performance comparison

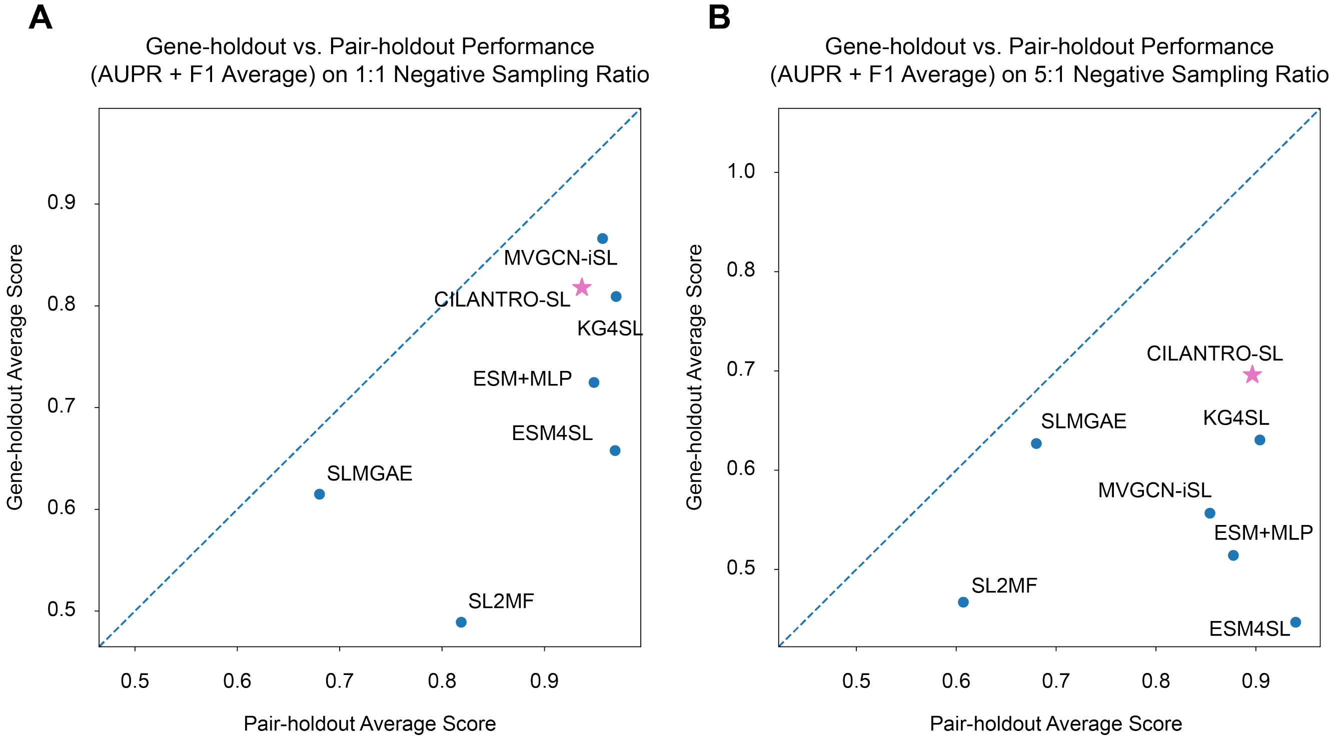


**Figure S8: Performance (average of AUPR and F1 Score) of models in different test-train split strategies.** We evaluated CILANTRO-SL against baseline models of KG4SL, $SL^2MF$, SLMGAE, MVGCN-ISL, ESM4SL, and ESM+MLP. Baseline DDGCN was not included as it does not take in negative samples, and has low AUPR and F1 scores (0.20518 and 0.37556 respectively). **A.** Performance on gene-holdout against pair-holdout in the 1x negative sampling setting. **B.** Performance on gene-holdout against pair-holdout in the 5x negative sampling setting.
